# A PHOX2B+ PONTINE NUCLEUS ESSENTIAL FOR INGESTION

**DOI:** 10.1101/2024.05.29.594519

**Authors:** Selvee Sungeelee, Caroline Mailhes-Hamon, Zoubida Chettouh, Philip Bokiniec, Annaliese Eymael, Bowen Dempsey, Jean-François Brunet

## Abstract

The first phase of feeding consists in acquiring solid foods from the environment by biting, and their preparation for swallowing by chewing. These actions require the precise coordination of tens of orofacial muscles for the jaw and tongue. The siege for this motor patterning is known to be in the reticular formation, a complex and poorly mapped region of the hindbrain, but the neuron groups involved are still elusive. Here, we characterize a group of reticular interneurons located in the supratrigeminal area that express the homeodomain transcription factor *Phox2b*. This nucleus — Sup5*^Phox2b^*— is premotor to both jaw-closing and jaw-opener motoneurons and receives direct input from cranial sensory afferents, motor cortex and satiation related nuclei. Its activity differentially tracks lapping, biting and chewing movements, suggesting its involvement in the elaboration of distinct orofacial motor patterns in vivo. Acute global activation or inhibition of Sup5*^Phox2b^*by optogenetics both interrupt volitional feeding sequences. Thus, Sup5*^Phox2b^*is an obligatory subcortical node, topologically and genetically defined, in the neural circuits that control the oral phase of feeding.

**Teaser:** A genetically defined cluster of neurons in the hindbrain is an essential relay for biting and chewing food.

## Introduction

Lapping, biting and chewing are the first phases of ingestion of food. They are executed by muscles of the jaw and tongue, controlled by two classes of cranial motor nuclei in the pons, medulla and cervical spinal cord: branchiomotor nuclei for the jaw and suprahyoid muscles — trigeminal (principal and accessory (Mo5 and Acc5)) and facial (principal and accessory (Mo7 and Acc7)); and somatomotor nuclei for the tongue and infrahyoid muscles —hypoglossal (Mo12) and an unnamed cervical motoneuron group (*1*) that we call MoC (*2*). Triggering and patterning the activities of Mo5, Mo7, Mo12 and MoC, i.e. mobilizing in a coordinated fashion the many muscles involved in lapping, biting and chewing, is thought to take place in the pons and/or medulla, as evidenced by the preservation of these complex feeding movements in decerebrate or deafferented animals (Miller and Sherrington, 1915; Woods, 1964)(*3–5*).

Like all motor circuits, those for orofacial feeding movements can be parsed, conceptually, into three elements upstream, i.e. premotor, to the motoneurons themselves: central rhythm generator (for those movements that are rhythmic, i.e. lapping and chewing); central pattern generator (that coordinate the muscles involved); and sensory inputs for adjusting the movements to a changing environment. In practice, the degree to which these circuit modules are distinct, and the hierarchical level that they occupy upstream of motoneurons are often speculative, the same neurons potentially subserving several functions (see below).

The central rhythm generators for chewing and lapping — which mobilize the same muscles for the most part, thus might be overlapping (discussed in (*6*)) — are still largely elusive. Potential locations— based on in vitro *en bloc* preparations (*7*) and pharmacological infusions of muscimol (*8*, *9*) or lidocaine (*10*) — are all in the reticular formation, mediolaterally encompassing its intermediate and parvocellular divisions, and rostro-caudally extending from the nTS to rostral Mo5 (reviewed in (*11*)). This relatively large brain region is also where many premotor neurons were mapped, most recently by monosynaptic retrograde tracing (*2*, *12–14*). These premotor neurons can be construed as part of the central pattern generator by virtue of their branching projections, which enable them to co-stimulate different muscles (*2*, *13*) or the same muscle bilaterally (*12*). The primary sensory neurons relevant for chewing are those of the periodontal receptors and jaw-closing muscle spindles, in the mesencephalic nucleus of the trigeminal nerve (Me5) and the trigeminal ganglion, the latter projecting onto second order sensory neurons in the trigeminal sensory nuclei (reviewed in (*15*)). First order sensory neurons in Me5 double as premotor neurons to Mo5 (*13*, *14*). Second order sensory neurons in the principal and oral part of the spinal trigeminal nuclei also have projections to Mo5 (reviewed in (*15*)) and display rhythmic patterns of activity during cortically-evoked chewing, proposed to reflect a role in central rhythm generation (*16*).

Demonstrating, *in vivo*, a causative role in orofacial feeding movements for any cell group of the reticular formation has been hampered, in recent years, by the dearth of genetic signatures, beyond those that underlie fast neurotransmitter phenotype. As a consequence, triggering, suppressing or modulating these movements has been so far mostly achieved by relatively unspecific or indirect manipulations of the PCRt: pharmacological interference with GABA_A_ receptors in the PCRt/IRt (*8*, *9*, *17*); optogenetic and chemogenetic manipulation of all GABAergic PCRt neurons (*18*); or optogenetic inhibition of inputs to the PCRt from the central amygdala (*18*)); in some cases movements were triggered together with a more holistic feeding behavior (*18*), or even with hunger-related depression of thermogenesis (*17*). To our knowledge, the most direct and specific manipulation of candidate medullary or pontine centers relevant for feeding was that of premotor neurons to Mo5 in the supratrigeminal nucleus — optogenetically accessed through their bilateral projections to Mo5, but irrespective of cell type— whose stimulation and inhibition both increased masseter tone, albeit without triggering or blocking jaw movements (*12*); and of IRt*^Phox2b^*, whose optogenetic excitation triggers lapping (*2*).

The landscape of neuronal types is best mapped by the expression of transcription factors, among which homeodomain proteins feature prominently (*19*, *20*). In the central nervous system, the expression of the homeoprotein Phox2b, is restricted to branchial motoneurons and groups of hindbrain interneurons. Here, we study a cluster of pontine *Phox2b-*positive interneurons that reside in the pons, at the location previously described as the supratrigeminal nucleus (Sup5), proposed to contain premotor neurons to Mo5 — a century ago by means of Golgi stains (Lorente De No, 1922), and most recently by monosynaptic retrograde tracing (Stanek et al., 2014a; Takatoh et al., 2021b). We show that *Phox2b* neurons of Sup5 (hereafter Sup5*^Phox2b^*) target Mo5, but also Mo12 and Mo7 and broad regions of the IRt/PCRt. In alert animals, both photo-stimulation and photo-inhibition of Sup5*^Phox2b^* totally block bouts of volitional feeding. Moreover, Sup5*^Phox2b^*activity, monitored by fiber photometry, differentially tracks lapping, biting and chewing, suggesting a role in fine-tuning the orofacial motor output according to feeding modalities.

## Results

### Topology and developmental origin of Sup5*^Phox2b^*

Mo5 is surrounded by the reticular formation, which in that region can be parsed into a peritrigeminal area (Peri5) ((*15*), first described as “regio h” in plate XII of (*21*)); and a supratrigeminal area (Sup5), first described and named on Figures 1-4 of (*22*). Both Peri5 and Sup5 contain many neurons that express the pan-visceral transcription factor *Phox2b* (*2*, *23*) (and **Fig. 1a**). For further study, we elected the supratrigeminal *Phox2b*^+^ population, hereafter referred to as Sup5*^Phox2b^*.

**Figure 1:**
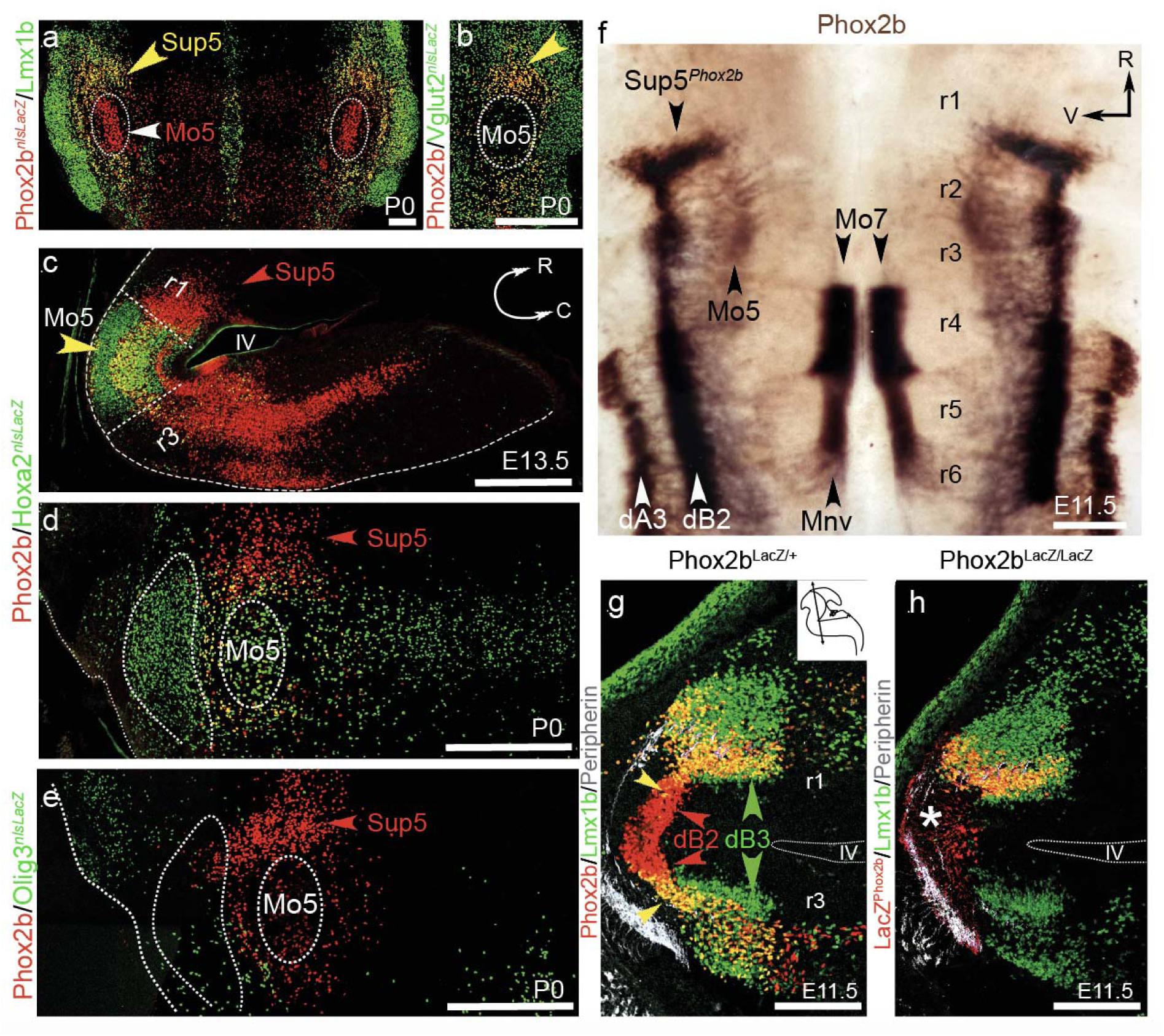
Genetic markers and ontology of Sup5 *^Phox2b^*. (**a**,**b**) Coronal sections at P0 through the pons of a *Phox2b::Cre;Tau-nlsLacZ* (**a**), and a *Vglut2::Cre;Tau-nlsLacZ* mouse (**b**), stained for the indicated markers. (**c**, **d**) Sections through the brainstem of a *Hoxa2::Cre;Tau-nlsLacZ mouse*, parasagittal at E13.5 (**c**, pontine curvature outlined in white) and coronal at P0 (**d**, principal trigeminal nucleus and Mo5 outlined by stippled line), immunostained for the indicated markers. (**e**) Coronal section through the brainstem of an *Olig3::cre^ERT2^;Tau nlsLacZ* mouse at P0, labeled with the indicated markers. (**f**) Flatmount of an E11.5 embryo hybridized with a *Phox2b* probe (previously analyzed in (*64*)). Precursors of Mo7 are still being produced from the pMNv domain of r4 and migrate caudally into r6. Precursors of Mo5 have already migrated dorsally in r2 and settled immediately ventral to the dB2 progeny. Sup5*^Phox2b^* neurons are born in the rostral-most pole of dB2, in r1. (**g**,**h**) Coronal sections (inset) through the pontine flexure of E11.5 embryos, heterozygous (**g**) or homozygous (**h**), for a null *Phox2b* allele, labeled with the indicated markers. The dotted outline indicates the recess of the 4th ventricle, above and below which, r1 and r3 are transversally sectioned, providing mirror images of the progeny of dB2 (Phox2b+) and of dB3 (unlabeled and giving rise to Lmx1b + postmitotic neurons (green arrowheads). The yellow arrowhead indicates the onset of Lmx1b expression in the dB2 progeny (red arrowheads) in r1 (i.e. the prospective Sup5*^Phox2b^*), as well as in r3. In the absence of *Phox2b* (**h**) the dB2 domain does not generate postmitotic neurons (white asterisk in **h**) but postmitotic *Lmx1b* neurons are still observed in r1. IV, fourth ventrical; C, caudal; Mo5, trigeminal motor nucleus; Mo7, facial motor nucleus; MnV, branchiovisceral motoneuronal precursors; R, rostral; r, rhombomere; Sup5, supratrigeminal nucleus; V, ventral. Scale bars: (**a**), 200 µm; (**b**): 500 µm; (**c**, **d**, **e**) 500 µm; (**f**), 400 µm; (**g, h**), 200µm.

**Figure 2:**
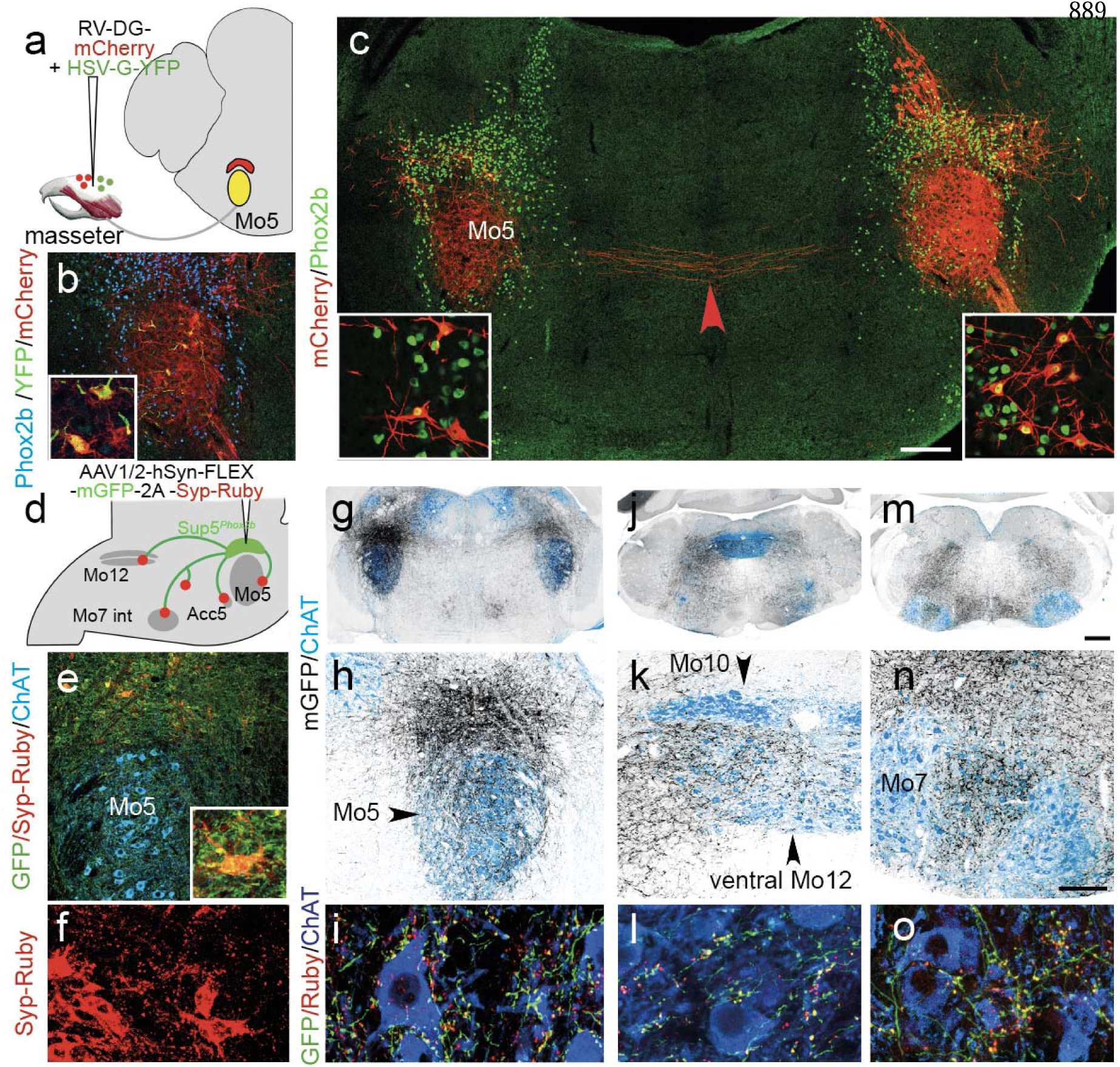
Synaptic targets of Sup5*^Phox2b^*. (**a-d**) Monosynaptically-restricted labeling of premotor neurons from the masseter in P8 wild-type pups. (**a**) Schematic of the strategy: G-deficient rabies virus (RV) expressing mCherry are co-injected into the masseter along with a YFP-expressing helper virus HSV-YFP-G, providing G-complementation. (**b**) Coronal section through the trigeminal region at P8 (higher magnification as inset), showing mCherry and YFP co-infected “seed cells” in Mo5 (yellow). (**c**) Coronal section through the pons showing premotor neurons (red) that express Phox2b (green) in ipsilateral and contralateral Sup5 (higher magnification as insets). Red arrowhead: commissural fibers. (**d**-**o**) Anterograde tracing from Sup5*^Phox2b^* neurons. (**d**) Schematic of the injection strategy with summary of the main projection sites. (**e**) Infected neurons in Sup5*^Phox2b^* express mGFP (green) and syp-Ruby (red). Inset: high magnification of an infected neuron in Sup5*^Phox2b^* itself covered with mGFP+/syp-Ruby+ boutons. (**f**) Dense syp-Ruby puncta from infected Phox2b+ cells in the Sup5 region itself. (**g**-**o**) Coronal sections through the brainstem at low (**g**, **j**, **m**), and higher (**h**, **k**, **n**) magnifications showing anterograde labeling (grey) of cranial motor nuclei (blue) from Sup5*^Phox2b^*. (**i**, **l**, **o)** Close-ups of motoneurons in the respective motor nuclei, showing axons (mGFP, green) and terminal boutons (syp-Ruby, red puncta) on the cell bodies. Scale bars: **g**, **j**, **m**, 500 µm; **b**, **c**, **e**, **h**, **k**, **n**, 200 µm; **i**, **l**, **o**, 20 µm.

**Figure 3:**
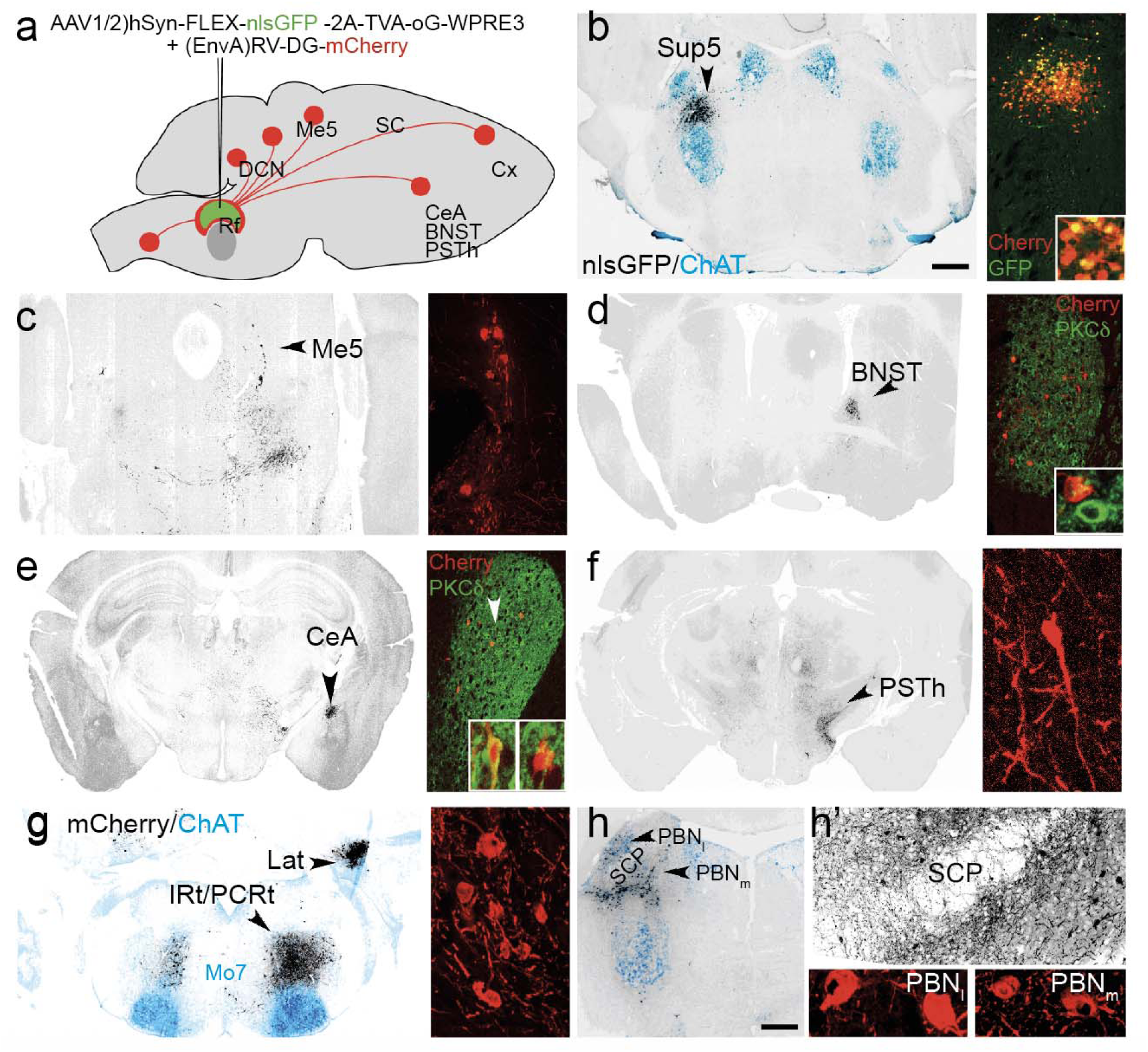
Inputs to Sup5^Phox2b^. (**a**) Strategy for monosynaptic retrograde tracing from Sup5*^Phox2b^*and summary of the major inputs. (**b**-**h’**) Low magnification (left panels) and higher magnification (right panels and insets) of coronal sections through the brain of adult mice showing cells that provide direct inputs to Sup5*^Phox2b^* (mCherry+ cells, grey in wide views, red in close ups). (**b**) The Sup5 region contains Sup5*^Phox2b^*seed cells (nlsGFP^+^/mCherry^+^) intermingled with input neurons (nlsGFP^—^/mCherry^+^). Dotted outilne denotes Mo5. (**c**) The first-order sensory neurons of the mesencephalic trigeminal nucleus (Me5) are retrogradely labeled from Sup5*^Phox2b^*. (**d**,**e**) The extended amygdala (i.e., the bed nucleus of the stria terminalis (BNST) (**d**) and the central amygdaloid nucleus (CeA) (**e**)), project to Sup5*^Phox2b^* ipsilaterally. Most of the retrogradely-filled neurons are PKC-∂^—^ (right inset) with an occasional positive one (white arrowhead and left inset); (**f**, **f’**) Posterior subthalamic nucleus (PSTh); (**g**) the Lateral dentate cerebellar nucleus (Lat); the same plane of section captures input from the ipsilateral intermediate (IRt) and parvocellular (PCRt) reticular formations and contralateral IRt; (**h**,**h’**) the parabrachial nucleus (PBN) (medial and lateral divisions). Other sites of input are in **Supplementary** Fig. 3. SCP: superior cerebellar peduncle. Scale bars: **b**-**h**, 500µm.

**Figure 4:**
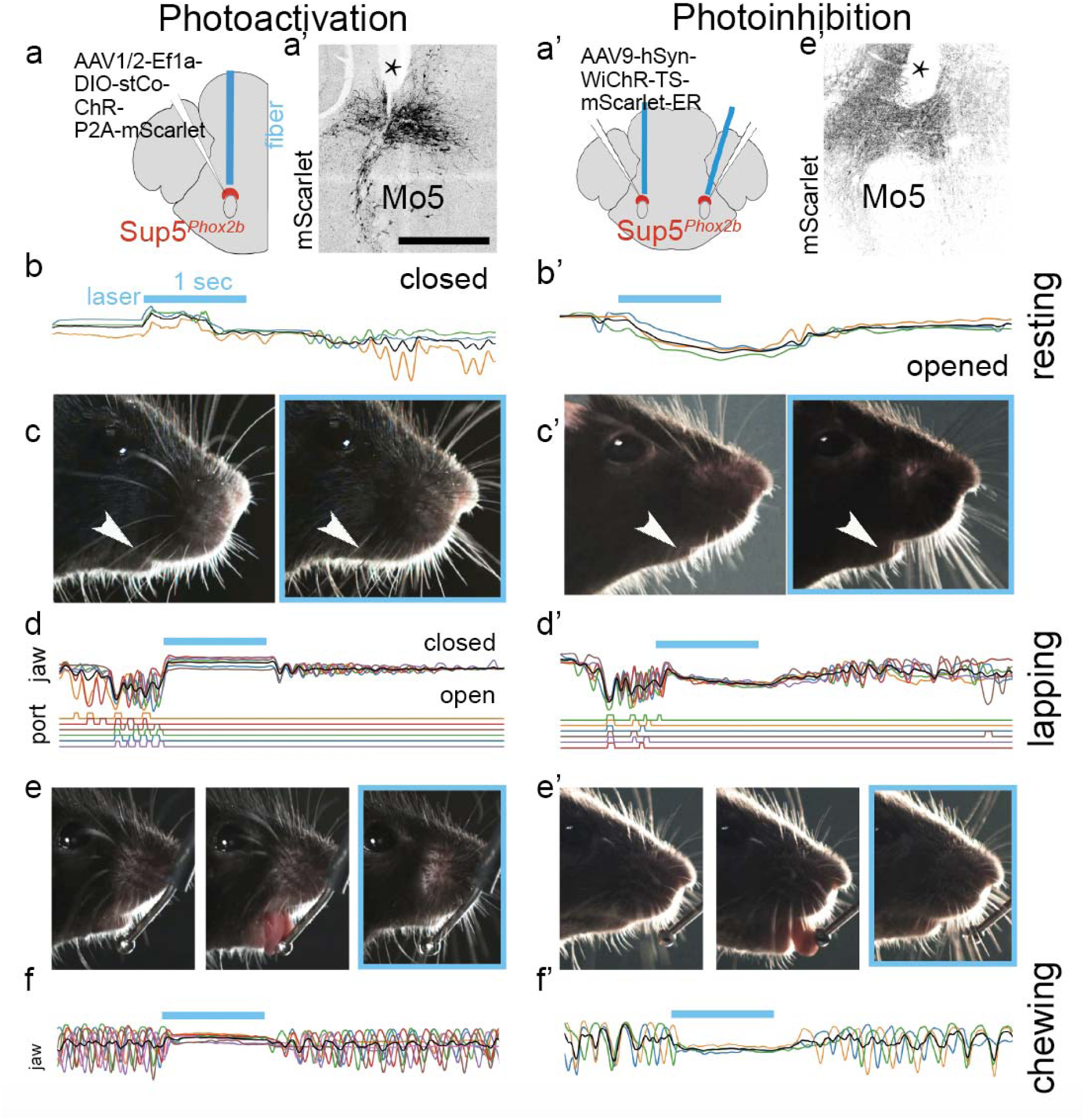
Global activation and inhibition of Sup5*^Phox2b^*. **(a-d’)** Photoactivation of Sup5*^Phox2b^*. (**a**,**a’**) Schematic of the viral injection and fiber-optic implantation for stimulation of Sup5*^Phox2b^* (**a**) and coronal section through the hindbrain showing infected Sup5*^Phox2b^* neurons and the optic fiber track in the brain (asterisk in **a’**). (**b-c**) Stacked traces of tracked jaw position on the Y-axis, during and after laser stimulation (blue bar) from one mouse (4 trials) (**b**) and example frames of the mouse’s face before and during stimulation (blue frame) (**c**). (**d,e**) Stacked traces of tracked jaw position on the Y-axis while lapping, from one mouse (6 trials), interrupted by the laser (blue bar) (**d**) and example frames of the mouse face during lapping (left frames) and during stimulation (blue frame) (**e**). (**f**) Stacked traces of tracked jaw position from one mouse while chewing (7 trials), interrupted by stimulation (blue bar). (**a’**,**f’**) equivalent schematic and data for photoinhibition. Scale bar: **a’**, **a’**, 500µm.

Sup5*^Phox2b^* caps Mo5 and lies medial to the dorsal aspect of the principal trigeminal nucleus (**Fig. 1a**). This nucleus co-expresses the homeobox transcription factors *Phox2b* and *Lmx1b* (**Fig. 1a**) and the vesicular glutamate transporter *VGlut2* like all surrounding reticular formation (**Fig. 1b**), but not the GABA transporter *GAD67*, which is expressed by other neurons intermingled with Sup5*^Phox2b^* (**fig. S1a**).

On the rostro-caudal axis of the hindbrain, Sup5*^Phox2b^*originates, and resides rostral to Mo5, in topological or genoarchitectonic terms(*24*). This is best evidenced in the *Hoxa2::Cre;Rosa^Tau-nlsLacZ^*genetic background, that labels derivatives of the second segment of the hindbrain (rhombomere 2) (*25*), where Mo5 is LacZ-positive (consistent with its origin mostly in rhombomere 2) while Sup5*^Phox2b^* resides immediately rostral, in a LacZ-negative region corresponding to rhombomere 1 (**Fig. 1c, d**). This topology is also manifested by the position of Sup5*^Phox2b^* rostral to Lbx1^+^ interneurons, thus to the boundary between rhombomeres 2 and 1 (*26*) (**fig. S1b**). Unlike Sup5*^Phox2b^*, most *Phox2b+* cells in Peri5 are derived from rhombomere 2 (**Fig. 1d**) (and **Fig. 2g** in (*2*)), including its rostral aspect, contiguous with Sup5*^Phox2b^*. Thus, Peri5*^Phox2b^* and Sup5*^Phox2b^*have distinct developmental origins.

We then explored the origin of Sup5*^Phox2b^* on the dorsoventral axis of the hindbrain. The *Lmx1b^+^/Phox2b^+^*signature displayed by Sup5*^Phox2b^*(**Fig. 1a**) was previously described as typical of the dorsal progenitor dA3 domain, from which the glutamatergic cells of the nucleus of the solitary tract are born (*27*, *28*) together with IRt*^Phox2b^* (*2*). However, Sup5*^Phox2b^* is not derived from dA3, as evidenced by lineage tracing in a mouse line where the *Cre* recombinase is under the control of *Olig3*, a determinant of dA3 neurons (**Fig. 1e**). Instead, as apparent on flatmounts of the hindbrain at E11.5 (**Fig. 1f**) and on sections through the pons at E12.5 (**Fig. 1g**), Sup5*^Phox2b^* appears to arise from dB2, whose progenitors express *Phox2b* (*27*) and — as shown in this study — switch on *Lmx1b* post-mitotically in r1-r3 (arrowheads in **Fig. 1g**), but not further caudally (**fig. S1b** and (*28*)). The *Phox2b^+^/Lmx1b^+^*postmitotic cell group, however, has an unusual appearance in that it extends ventrally from the dB2 domain and is entirely contained within a wider population of *Phox2b*^—^/*Lmx1b^+^* cells in r1 (**Fig. 1f, g**). This pattern looks ambiguously like it could result from the postmitotic onset of *Lmx1b* expression in the *Phox2b*+ progeny of dB2, or the postmitotic onset of *Phox2b* in a *Lmx1b+* population arising from the ventrally abutting dB3 domain (*27*, *28*). An origin in dB2 is supported by **Fig. 1g** (arrowheads); an origin in dB3 is supported by *Phox2b* KO embryos where dB2 progenitors fail to produce post-mitotic cells (red arrowhead in **Fig. 1h**) and remain in the state of radial glia (white asterisk), but where an incipient Sup5*^Phox2b^* was present nevertheless in r1 (**Fig. 1h**). All in all, despite its anatomical coherence and unifying *Phox2b/Lmx1b* signature, Sup5*^Phox2b^* might have a dual dB2/dB3 origin (thus a possible heterogeneity) that we could not resolve with available tools.

Another ontological landmark of Sup5*^Phox2b^* is that it aggregates, from its inception, around the mesencephalic tract of the trigeminal nerve, i.e. the projections from the mesencephalic nucleus of the trigeminal nerve (Me5) (**Supplementary** Fig.1), which will constitute one of its major sources of inputs (see below).

### Connectivity of Sup5*^Phox2b^*

The original definition of Sup5 — poorly differentiated cytoarchitecturally from the principal trigeminal nucleus on its lateral side, and from the reticular formation on its medial side (discussed in (*29*))— was mostly based on connectivity: a major site of projections from Me5, and of inputs to Mo5 (*22*). We asked whether these hodological features hold true for Sup5*^Phox2b^*.

First, we assessed whether Sup5*^Phox2b^* is premotor to Mo5 by monosynaptic retrograde tracing from the masseter. In the masseter of pups at postnatal day 3 (P3), we injected a modified G-deficient rabies virus expressing mCherry together with a G-complementing helper virus expressing a YTB reporter, and analyzed brains at P8 (**Fig. 2a**). Seed cells co-expressing mCherry and YTB were in the dorsolateral Mo5 ipsilaterally, consistent with the known somatotopic representation of jaw closers in this nucleus (*30*) (**Fig. 2b**). Monosynaptically-traced cells resided in lateral Sup5, both on the injection and contralateral sides (**Fig. 2c** and insets), most of them expressing *Phox2b*. (They were the only Mo5 premotor neurons expressing the gene). Therefore, at least a subset of cells in Sup5*^Phox2b^*are premotor to jaw-closing motoneurons in Mo5. This is in line with previous observations that Sup5 cells target Mo5 in transgenic *Phox2b*-GFP rats (*23*). The lateral bias in the distribution of Sup5*^Phox2b^* premotor neurons could reflect the somatotopy of the injection in the masseter (which is made of several fascicles), or the early developmental phase at which the experiment was done (before weaning, and therefore presumably before the full engagement of the masseter in feeding).

Next, we evaluated the brain-wide projections of Sup5*^Phox2b^*by injecting it with a Cre-dependent AAV construct that co-expresses a membrane-tethered GFP and a reporter-tagged synaptophysin (Syp-Ruby) to label synaptic boutons (*31*) (**Fig. 2d**). The most proximal site of pre-synapses from Sup5*^Phox2b^*was coextensive with Sup5*^Phox2b^* (**Fig. 2 e,f**). At least some of these synapses were made on Sup5*^Phox2b^* cells themselves, as evidenced by boutons on the primary infected cells (inset in **Fig. 2e**). As expected from the retrograde tracing from the masseter, the second densest site of projections was Mo5, containing jaw adductors (**Fig. 2g-i**). In line with the necessity to retract the tongue while closing the jaw, Mo12 was also targeted, preferentially in its retrusor compartment (**Fig. 2j-l**). Thus, like IRt*^Phox2b^*(*2*) and other groups of orofacial pre-motoneurons (*13*, *32*), Sup5*^Phox2b^*is in a position to coordinate several orofacial muscles expected to synchronize. Less expectedly however, Sup5*^Phox2b^* also targeted, albeit at lower density than Mo5, its antagonists (jaw-opener motoneurons): the accessory 5th nucleus (Acc5) (**fig. S2a**), intermediate Mo7 and accessory 7^th^ nucleus (Acc7) (**Fig. 2m-o**) and MoC (**fig. S2b, b’**). Projections and boutons also covered the salivary preganglionic neurons dorsal to Mo7 (**fig. S2c-c”**), a possible substrate for the masticatory-salivary reflex (*33*). All motor nuclei were targeted bilaterally with ipsilateral predominance.

Other sites were, rostral to Sup5*^Phox2b^*, the CeA and BNST (**fig. S2d, e**), the ventral posteromedial thalamic nucleus (**fig. S2f**), the deep mesencephalic nucleus (**fig. S2g**), and caudal to Sup5*^Phox2b^*, the pontine reticular nucleus, caudal part (**fig. S2h, h’**) and broad regions of the caudal medullary reticular formation (**Fig. 2j**). The remarkably wide range of projections of Sup5*^Phox2b^* suggests either that this putative premotor center controls motoneurons both directly and indirectly, or that it has a more integrative role in feeding behaviours.

Next, we identified the source of inputs to Sup5*^Phox2b^*by injecting Sup5 of *Phox2b::Cre* mice, first with a *cre*-dependent AAV construct expressing the optimized rabies glycoprotein oG, TVA receptor and nuclear GFP; then with an EnvA-pseudotyped G-deficient rabies virus encoding the fluorophore mRuby (**Fig. 3a**). Neurons in Sup5*^Phox2b^* that co-expressed both vectors (i.e. seed cells) were thus doubly labelled with GFP and mCherry (**Fig. 3a, b**), while their presynaptic partners were single-positive for mCherry. Again, the most proximal of the latter were inside Sup5*^Phox2b^* itself (**Fig. 3b** and inset), comprising either other Sup5*^Phox2b^*cells (see above), or cells of another type, yet spatially contained within Sup5*^Phox2b^*. Thus, the combined anterograde and retrograde tracings show that Sup5*^Phox2b^* is bidirectionally connected within its own confines, at least in part with other Phox2b cells, although we could not ascertain the identity of all synaptic partners. The second most proximal source of inputs was, as expected from (*22*), the ipsilateral trigeminal mesencephalic nucleus (**Fig. 3c**). Other sites of input were, from rostral to caudal: the primary motor and insular cortices (Cx) (**fig. S3a, b**), the extended amygdala (CeA and BNST) (**Fig.3d**,**e**), the parasubthalamic nucleus (PSTh) (**Fig. 3f**), the deep (lateral) superior colliculus (**fig. S3c**), the lateral (**Fig. 3g**) and other deep cerebellar nuclei (**fig. S3d**), the parabrachial nuclei (**Fig.3h**, **h’**), the IRt and PCRt (**Fig.3g, fig. S3e**).

### Global stimulation or inhibition of Sup5*^Phox2b^* prevent volitional lapping and chewing

To investigate the role of Sup5*^Phox2b^* in vivo, we injected a *Cre*-dependent opsin (AAV1/2-Ef1a-DIO-stCoChR-P2A-mScarlet) (*34*) into the Sup5 of *Phox2b::Cre* mice and unilaterally implanted an optic fiber above this nucleus (**Fig. 4a**). We then optogenetically stimulated these cells with 1000 ms single pulses in head-fixed mice and video-recorded the face at rest ((**Fig. 4b, b’**) and **Supplementary video 1**). Optogenetic stimulation triggered an abrupt jaw adduction (of small amplitude since it occurred from the resting, essentially closed, position) in all mice (n=4), followed by return to the baseline position after about 700 ms. Photostimulation of Sup5*^Phox2b^* with 100 ms pulses at 5Hz evoked repetitive jaw adduction from the resting position at the same frequency (**Supplementary Video 2**), showing that Sup5*^Phox2b^* can operate within this frequency range. Thus, Sup5*^Phox2b^* can close the jaw, which is coherent with its projections to jaw closer motoneurons in Mo5. We then asked whether tonic photostimulation of Sup5*^Phox2b^* could disrupt volitional ingestive sequences. We established an optogenetic stimulation protocol during lapping in head-fixed mice, whereby 4 successive licks trigger a 1000 ms optogenetic stimulation of Sup5*^Phox2b^*. During this protocol, all lapping activity was abolished concurrent with jaw adduction ( **Fig. 4c, c’**, **and Supplementary video 3).** Small vertical and horizontal jaw movements occurred immediately after optogenetic stimulation, although lapping per se took longer to resume. This sequence was consistent across all tested animals (n=4). Similarly, photostimulation of Sup5*^Phox2b^* during chewing of a flake of almond consistently interrupted the chewing sequence (**Fig. 4d, Supplementary video 4**).

To investigate whether acutely silencing Sup5*^Phox2b^*impacts orofacial ingestive movements, we bilaterally injected WichR1 (a K^+^-selective channelrhodopsin (*35*)), into the Sup5 region of *Phox2b::Cre* mice (**Fig. 4e**) and photo-inhibited Sup5*^Phox2b^*tonically for 1 second, during periods of rest, lapping, or chewing in a head-fixed position. At rest, photoinhibition elicited a slight abduction of the jaw from its resting position (suggesting relaxation of jaw adductors), that persisted throughout the stimulation period, and even beyond (**Fig. 4f, f’, Supplementary video 5**). During either lapping (**Fig. 4g, g’** and **Supplementary video 6)** or chewing (**Fig. 4h** and **Supplementary video 7**), photoinhibition terminated the ingestive sequence, immobilizing the jaw for the length of the laser signal in a position slightly abducted compared to that caused by photostimulation. Random jaw movements resumed immediately after the end of the laser signal, but it took much longer and variable intervals for the animal, possibly disturbed by the interruption, to start eating again.

Thus, volitional ingestion of liquids or solids cannot proceed upon the global activation or inhibition of Sup5*^Phox2b^* neurons.

### The activity of Sup5*^Phox2b^* tracks ingestive orofacial movements

Finally, we tested whether Sup5*^Phox2b^* is active during spontaneous ingestive sequences. We carried out in vivo bulk fluorescent calcium recordings of Sup5*^Phox2b^*during spontaneous lapping and chewing in head-fixed mice using fiber photometry(*36*). We injected a Cre-dependent GCaMP7s vector (*37*) into the Sup5 of *Phox2b::Cre* mice and an optical fiber was implanted unilaterally above this nucleus to measure changes in Ca2+ dynamics while monitoring jaw movements (**Fig. 5a**). In all tested animals (n=3), lapping bouts correlated with increases in fluorescence of Sup5*^Phox2b^*(**fig. S4a**), indicating that Sup5*^Phox2b^* is recruited during lapping. When mice ate an almond flake (**Fig. 5b),** Sup5*^Phox2b^*was also recruited and its activity was much increased (about 5 fold) compared to lapping (**fig. S4a**). Moreover, the calcium changes in Sup5*^Phox2b^*tracked the microdynamics of almond ingestion: Gcamp7s fluorescence peaked with each biting event, followed by a slow decay of calcium fluorescence while chewing (**Fig. 5b**), during which smaller peaks correlated with each jaw movement (frequency = 5 to 6Hz) (**Fig. 5b, fig. S4b, Supplementary video 8**). These two nested patterns (reproducible for n=3 animals, across n=3 chewing sequences) suggest either a switch from tonic to rhythmic activity of Sup5*^Phox2b^*, or the successive operation of two neuronal populations within Sup5*^Phox2b^*, underlying, respectively, the occasional events of biting and the rhythmic movements of chewing in between. Switching, during the same trial, from almond to raw pasta (tougher and less brittle than almond) (**Fig. 5c**), led to a much higher frequency of biting events and a shortening of chewing sequences, resulting in a more constant activity of Sup5*^Phox2b^* and the transition from a cyclic alternation of biting and chewing, to a more chaotic pattern (**Fig. 5c**).

**Figure 5:**
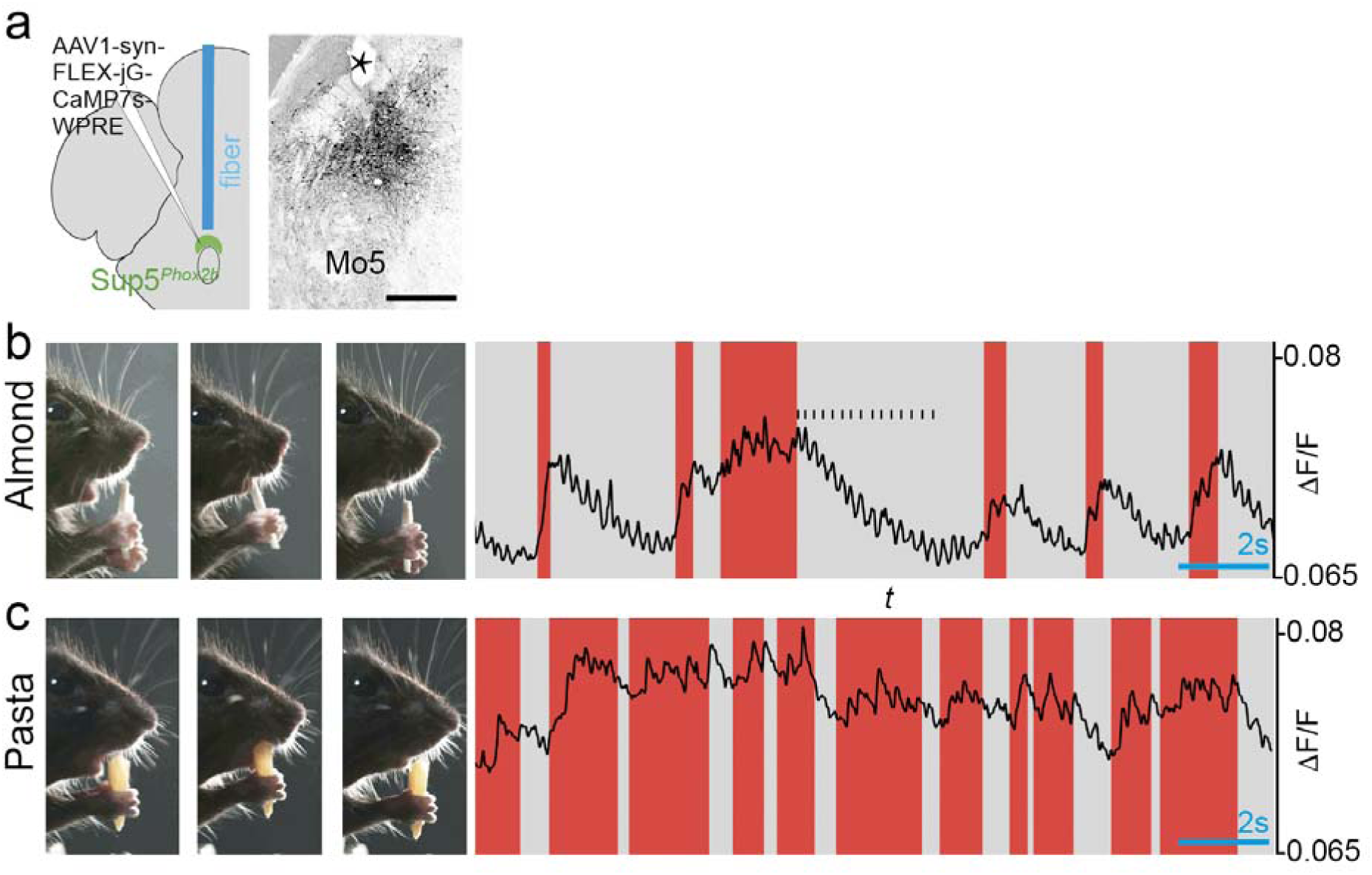
Activity of Sup5*^Phox2b^* during biting and chewing. (**a**) Schematic of the viral injection and fiber-optic implantation for fiber photometric recording of Sup5*^Phox2b^*, and transverse section through the hindbrain showing infected Sup5*^Phox2b^*neurons and the optic fiber track in the brain (asterisk). (**b**,**c**) Example frames of a mouse face at different phases of biting and chewing a flake of almond (**b**), or a piece of raw pasta (**c**): preparing (left), biting (middle), chewing (right) and example traces showing changes in bulk fluorescence (ΔF/F) of Sup5*^Phox2b^* during a recording session (∼18 sec) of eating an almond (**b**), or pasta (**c**), including bouts of biting/gnawing (red areas) and chewing episodes (grey areas). Rhythmic increases of fluorescence parallel each chewing movement of the jaw (black vertical marks). Scale bar: **a**, 500µm.

All in all, fiber photometry recording of Sup5*^Phox2b^* revealed that it differentially tracks orofacial movements: minimum in lapping, maximum in biting, and progressively waning during chewing while still tracking each chewing cycle, in a way that parallels the load force (minimal or null in lapping, highest when biting hard material, and decreasing during a chewing sequence, as the food is broken down).

## Discussion

We have identified Sup5*^Phox2b^*, a group of glutamatergic neurons marked by the transcription factor *Phox2b*, whose activity tracks jaw movement associated with lapping, biting and chewing in the alert animal, and whose global optogenetic perturbation, whether by stimulation or inhibition, blocks all spontaneous ingestive movements. Thus, a minimal functional attribute of Sup5*^Phox2b^* is that of an essential node in the premotor command for food and liquid ingestion. Sup5*^Phox2b^*could, for example, be an obligatory relay downstream of a central rhythm generator. One candidate is IRt*^Phox2b^*, provided that the IRt cells that send massive input to Sup5*^Phox2b^* (**Fig. 3g** and **fig. S3e**)) express *Phox2b*, thus belong to IRt*^Phox2b^* (which we could not ascertain for technical reasons). Indeed IRt*^Phox2b^* acts as a lapping central rhythm generator upon photo-stimulation (*2*); and since lapping involves many of the same muscles as chewing, coordinated in similar fashion, they have been proposed to share a central rhythm generator (*6*). Thus, IRt*^Phox2b^* could be the common central rhythm generator, downstream of which Sup5*^Phox2b^* implements a chewing/biting pattern, that would be shaped by the input from the jaw proprioceptors in Me5 (*15*)(**Fig. 3c**); or, by virtue of its reciprocal connections with IRt (**Fig. 3g** and **fig. S3e**), partake in rhythm generation by providing cyclical feedback to IRt*^Phox2b^*.The fact that Sup5*^Phox2b^* produced no rhythmic jaw movements upon photoactivation excludes *prima facie* the possibility that it is, by itself, a rhythm generator. However, this cannot be excluded at this point. Indeed photogenetic activation of a genetically-defined candidate central rhythm generator might be an intrinsically poor means of triggering rhythmic activity, and we are not aware of any published case so far — except, as it were, for IRt*^Phox2b^* and lapping (*2*): it could be that, in many excitation paradigms, the tonic bulk photoactivation of all potentially rhythmic cells prevents their rhythmic oscillations. Arguing that Sup5*^Phox2b^* should not be written off as an orofacial central rhythm generator, or part thereof, are that: i) it resides in the minimal block of tissue that can produce, *in vitro*, a 4-8Hz rhythm in the trigeminal nerve, or attached masseter and digastric muscles: “between the caudal pole of Mo5 and within 1mm rostral to Mo5” (*7*, *38*); ii) it receives and sends many connections within its own confines, suggesting the existence of a micro-circuitry, amplifying or self-limiting, of a type classically invoked to underlie pacemaker networks in vertebrates (*39*, *40*).

Other afferents to Sup5*^Phox2b^* place it in the position to integrate influences from higher centers, a striking proportion of which are associated with regulation of food intake, and which more precisely are anorexigenic: the central amygdala (*41*), the parasubthalamic nucleus (*42–44*), the bed nucleus of the stria terminalis ((*45*) and references within) and the lateral deep cerebellar nucleus (*46*). In turn, several of these sites are targets of Sup5*^Phox2b^* (CeA and BNST) (**Supplementary** Fig.2, **Fig. 3 and fig. S3**). It is not straightforward to predict the combined effect of these inputs, based on their known properties: the CeA cells that are presynaptic to Sup5*^Phox2b^* are presumably GABAergic (*47*), which could fit with an inhibitory role in ingestive behaviors, but they are for the most part PKC-∂^—^ (**Fig. 3**), i.e. are not the PKC-∂^+^ cells most clearly implicated in anorexia (*41*, *48*); on the other hand the PSTh is considered glutamatergic (*49*)), thus should activate Sup5*^Phox2b^*, although global stimulation could lead to functional suppression, similar to the effect we obtained with optogenetics. Whatever the integrated effects of these inputs, one can predict from the literature that it is negative; and one can hypothesize that it is preferentially on biting, i.e. on the initiation of food intake, rather than chewing, a fairly stereotypical action whose only useful modulation should come from oral sensations. In this context, it is notable that the activity of Sup5*^Phox2b^*, as measured by fiber photometry, is maximum during biting (**Fig. 5c**). Even though, in the context of feeding, mastication movements do not make sense, indeed never occur before food has entered the mouth, a recruitment of the premotor neurons by higher orexigenic centers (e.g. in a predatory context as in (*18*)), or their inhibition by anorexigenic centers (as suggested by the connectome of Sup5*^Phox2b^*) can be rationalized as the “priming” or “gating”, respectively, of an activity before it is required (*50*).

The expression of *Phox2b* in a lapping center (*2*), as well as an orofacial premotor center essential for ingestion (this study), expands the physiological correlate of this developmental transcription factor. *Phox2b* was originally discovered as a master gene for the “autonomic nervous system” as defined by Langley (*51*), i.e. for the autonomic outflow: preganglionic and ganglionic neurons of the sympathetic, parasympathetic and enteric nervous systems (with the exception of sympathetic preganglionics) (*52*); It later emerged that the developmental role of *Phox2b* correlates with a more holistic “visceral nervous system” (*53*) that includes the sensory afferents to this autonomic outflow (cranial visceral sensory neurons and their nucleus of projection, the nucleus of the solitary tract) (*54*); as well as the branchiomotor (or “special visceromotor”) neurons (*55*) to orofacial muscles, whose ancestral role was exclusively in visceral functions: feeding and breathing. Our characterization of IRt*^Phox2b^* and Peri5*^Atoh1^* (*2*) and our current analysis of Sup5*^Phox2b^*, reveal that upstream of *Phox2b*^+^ branchiomotor neurons, yet another layer of *Phox2b* neurons is involved in triggering and patterning their activity during lapping, biting and chewing (and presumably suckling). Thus, the landscape of *Phox2b* neurons, while transcending classical neuroanatomical entities (and crossing the border between “autonomic” and voluntary ones), outlines a novel ensemble, strikingly coherent from a physiological standpoint: the circuits that control all nutritive actions, from the capture of nutrients or fluids from the environment to the rejection of waste, through appetitive and aversive taste perception, salivation or other digestive secretions, and movements of the digestive tube. Somehow, it was evolutionarily expedient to recruit, over and over again, the same neural transcription determinant, *Phox2b*, for elaborating the outer and inner behaviors that provide the cells of the body with fluids and calories. The functional survey of *Phox2b* neurons is nearing completion but some remain to be investigated, e.g. derivatives of the dB2 domain caudal to r2, and thus to Mo5, in the parvocellular reticular formation (**Fig. 1c and fig. S1b,c**). It is an intriguing possibility that they will also partake in such behaviors.

## Materials and Methods

### Mouse lines

The following transgenic mouse lines were used in this study: *Phox2b::Cre* (*56*), *vGlut2::Cre* (*57*), *Tau::Syp-GFP-nlsLacz* (also known as *Tau-mGFP)*(*58*)*, HoxA2::Cre* (*25*) *Phox2b:LacZ* (*59*), *Olig3::CreERT2*L(*28*). All mouse lines were bred on a B6D2 F2 background. The experiments were performed on embryos at embryonic (E) days E9.5–17.5, neonate pups at postnatal day 2–8 (P1–8), and adult (P30–56) animals of either sex. All experimental procedures and protocols were approved by the Ethical Committee CEEA-005 Charles Darwin (authorization 26763-2020022718161012) and conducted under EU Directive 2010/63/EU. All possible measures were taken to minimize the suffering and number of animals used. The sequence of all primers for genotyping are in **Table 1**.

**Table 1:**
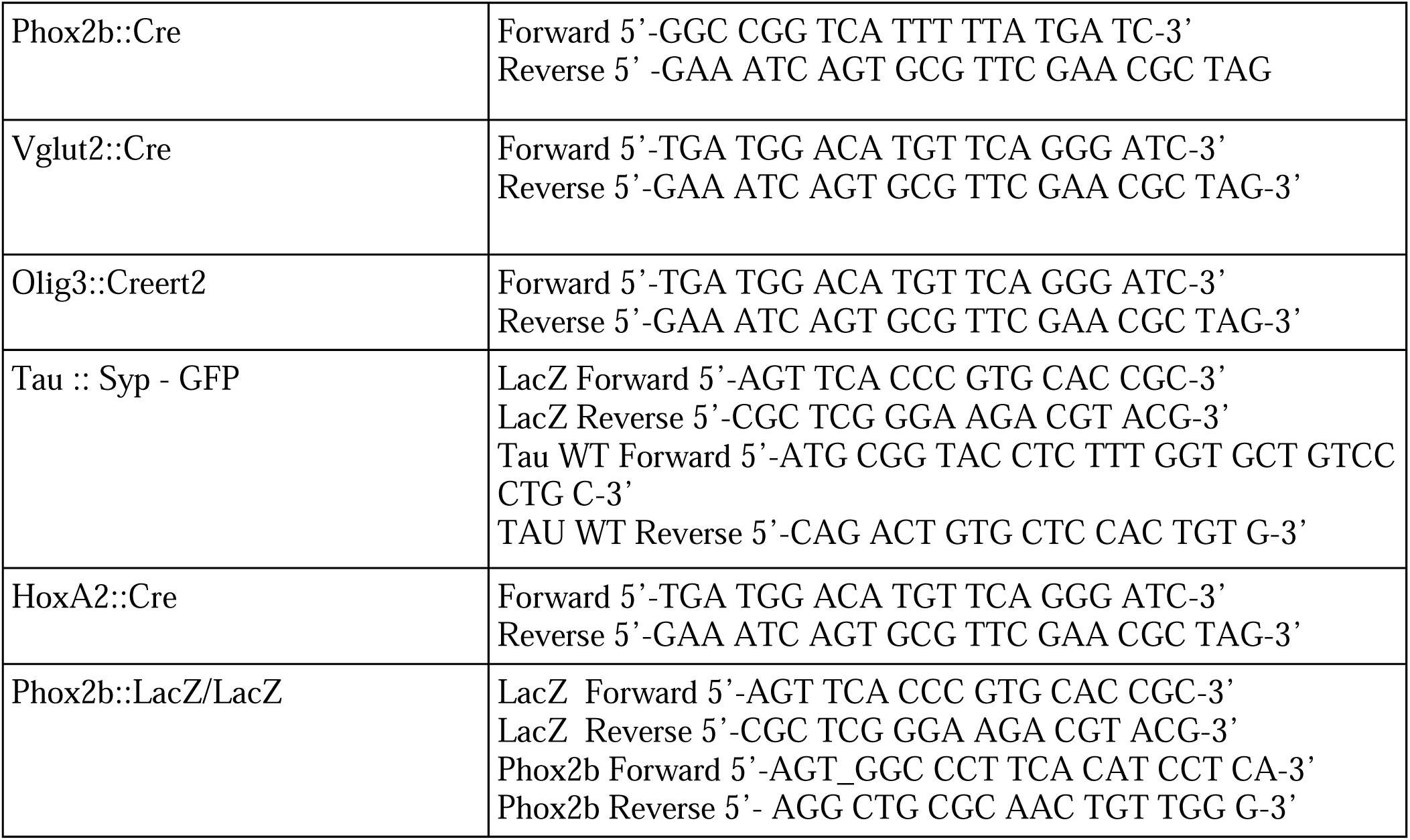
sequences of oligonucleotides used for genotyping.

### Animal husbandry and housing

Mice were housed in groups in conventional housing conditions, at 21-22°C, 40% to 50% humidity, with unlimited access to standard chow and water and kept in a 12-12-hour light/dark cycle.

Mice that underwent surgical implantation for optogenetics and photometry experiments were subjected to a 12-hour reverse light/dark cycle and tested in the dark phase.

### Tamoxifen injections and cre recombination

Cre-recombination in *Olig3-CreERT2* crosses at E9.5 was induced by intra-peritoneal injections of 50mg/kg body weight of tamoxifen in corn oil in pregnant dams.

### Viral vectors for tracing, optogenetic, and photometry experiments

Anterograde tracing from Sup5*^Phox2b^* was carried out by unilateral injection, 3-4 months old *Phox2b::Cre* mice, of 250Lnl of a Cre-dependent vector (pAAV-hSyn-FLEX-mGFP-2A-Synaptophysin-mRuby, titer: 7×10¹² viral genomes (vg)/ml, Addgene #71760-AAV1) which expresses membranal GFP in neurons and a red fluorescent reporter mRuby fused to the synaptophysin gene, which accumulates in the synaptic boutons, thereby outlining their synaptic targets (*31*)

To map the inputs to the masseter muscle, 50 to 100Lnl of a 1:1 viral cocktail of RV-B19-ΔG-mCherry (titer: 1.3L×L10^9^ TU/ml, Viral Vector Core—Salk Institute for Biological Studies) and an HSV-hCMV-YFP-TVA-B19G (titer: 3L×L108LTU/ml, Viral Core MIT McGovern Institute) was unilaterally injected into the masseter of mouse pups (P3).

To label inputs to Sup5*^Phox2b^* a two-step tracing strategy was used (*60*, *61*): first, 50 nl of a Cre-dependent AAV1/2-Syn-flex-nGToG-WPRE3 (titer: 8.1 × 1011vg/ml, Viral Core Facility Charité) was injected into the Sup5 of *Phox2b::Cre* mice. This vector expresses the genes for an EnvA-interacting receptor (TVA), a nuclear green fluorescent protein (nls-GFP), and an optimized G protein (oG)(*60*). Two weeks later, 100nl of EnvA-pseudotyped rabies virus (EnvA-RABV-SADB19-ΔG-mCherry; titer: 3.1 × 108 vg/ml, Viral Vector Core, Salk Institute for Biological Studies) was injected at the same coordinates. The rabies virus specifically infects AAV-containing, thus TVA+ neurons (seed cells), which subsequently enable its transport in presynaptic neurons by their expression of oG. Thus, seed cells express both nls-GFP and m-Cherry, presynaptic cells express m-Cherry alone. After an additional 7 days, the brains were harvested and the presynaptic neurons were identified based on their location in relation to the Paxinos Atlas (*62*).

For photoactivation experiments, Sup5*^Phox2b^* of *Phox2b::Cre* mice was unilaterally infected by AAV1/2-Ef1a-DIO-stCoChR-P2A-mScarlet (250Lnl, titer: 3L×L1013Lvg/ml, kind gift from O. Yizhar) that drives expression of a soma-targeted opsin.

Acute silencing of Sup5*^Phox2b^* was achieved by bilateral infection with of AAV9-hSyn-WiChR-TS-mScarlet-ER (250nl, titer: 0.8 x 1012vg/ml, Viral Core Facility Charité)) that encodes a K+-selective soma-targeted opsin.

For photometry experiments, a unilateral 250Lnl injection of AAV1-syn-FLEX-jGCaMP7s-WPRE (titer: 1L×L1012Lvg/ml, Addgene #104487-AAV1) was used to express GCAMP7s into Sup5*^Phox2b^*.

## Surgical procedures

### Stereotaxic injections and implants

*Phox2b::Cre* mice were anesthetized with an intraperitoneal injection of 50 mg/kg Zoletil (Zoletil 100, Virbac Sante Animale, France) and 10 mg/kg Xylazine (Rompun, 2%)).Thirty minutes before the start of surgery, buprenorphine ((0.3 mg/mL, 0.1mg per kg body weight, Buprecare)) was administered subcutaneously as analgesia. Mice’s core temperature was maintained within the physiological range using a homeothermic pad (37.5-38°C). Briefly, anesthetized animals were placed in a stereotaxic frame (Kopf), and 100μl of lidocaine (2%) was injected under the skin overlying the skull for local analgesia. To target Sup5*^Phox2b^*neurons, standard surgery was performed to expose the brain surface above Sup5 at the following stereotaxic coordinates: bregma −5.00mm, lateral ±1.40mm, dura –2.80mm.

For viral injections, 50-250 nL volumes were delivered at the rate of 75-100nL per minute via quartz glass capillary needles (QF100-50-7,5, WPI) backfilled with mineral oil with a 10μL Hamilton syringe (701 RN) connected to a pump (Legato 130, KD Scientific, Phymep, France). After infusion, the injection needle was maintained in position for 5 min to reduce backflow of the virus during needle retraction. For optogenetics experiments, 200 μm core optic fibers (0.39NA, Smart Laser Co., Ltd) were placed at the same dorsoventral coordinates as the injections. For photometry experiments, optic fibers (0.57 NA, Smart Laser Co., Ltd) were implanted 200 μm below the injection sites (Dura –3.00 mm). An anchor screw was then placed into the cranium anterior to the brainstem. The optic fibers were subsequently secured into the brain by UV-cured dental adhesive cement (Tetric Evoflow, Ivoclar Vivadent)applied to the skull, ceramic ferrule and anchor screw. Custom-made head-fixation implants (Ymetry, Paris France) were positioned onto the skull and similarly affixed with dental adhesive cement. For inhibition experiments, that require bilateral optic-fiber placement, one of the optic fibers was implanted at an angle of 5° above Sup5 (axis of rotation at ±1.40mm lateral from Bregma) to leave space for the head-bar on the skull between the two implants. The angled coordinates for the placement of the optic fiber to target Sup5*^Phox2b^*, calculated by basic trigonometry, were: bregma −5.00mm, lateral ±1.73mm, skull –3.81 mm.

Mice recovered from anesthesia on a heating pad for a day before being placed back into their home cages. Appropriate postsurgical care was provided, and animals were regularly monitored for signs of infection, pain, or lethargy until behavioral assays began.

### Intramuscular masseter injections

Masseter muscle injections were performed at P3 neonatal stage. Pups were anesthetized by deep hypothermia. For anesthesia induction, pups were placed in latex sleeves and gently submerged in crushed ice for 3-5 minutes. Anesthesia was maintained (up to 10 min) by placing pups on a cold pack (3–4L°C). A small incision was made in the skin overlaying the masseter and a glass pipette (tip diameter ca. 0.1 mm), filled with the virus cocktail and connected to a pneumatic dispenser (Picospritzer), was guided into the masseter with a 3D micromanipulator. Muscular filling with the viral cocktail was achieved with 5–10 pressure pulses (100 ms, 3–5 bars), which delivered around 50nL of the virus. This was confirmed by the spreading of Fast-Green dye (0.025%) added to the viral solution. After injection, the pipette was withdrawn and the incision was irrigated with physiological saline before suturing (10-0 gage suture (Ethilon)). Pups were promptly returned to the mother after a brief recovery onto a heating pad. The brains were harvested five days after injection (P8) for histology.

## Histology

### Immunofluorescence

Depending on the stage, histology of the brain was carried out either on whole embryos (up to E16.5) dissected out of uterine horns or on brains dissected out of the cranial vault for embryos E17.5 to P0. Adults and postnatal animals were euthanized by an intraperitoneal injection of pentobarbital (Euthasol Vet, 140mg/kg), transcardially perfused with ice-cold PBS followed by 4% PFA in PBS (Antigenfix, Diapath), and the brains were rapidly dissected.

Brains or embryos were then postfixed in 4% PFA overnight at 4 °C, rinsed 3 times for 30 minutes each in PBS then cryoprotected in 15% sucrose in PBS, overnight at 4°C. Tissues were subsequently embedded in gelatin-sucrose medium (7.5% gelatin in 15% sucrose in PBS) and frozen for cryo-sectioning at 30-60 μm on a CM3050s cryostat (Leica). Sections were washed for 1 h in PBS and incubated in a blocking solution (10% calf serum in 0.5% Triton-X100 PBS) containing the primary antibody, which was applied to the surface of each slide (300 μL per slide). Slides were placed in a humidified chamber on a rotating platform and incubation lasted 4– 8 h at room temperature and overnight at 4°C. Sections were washed in PBS (3 × 10 min), then incubated in the dark with the secondary antibody in a blocking solution for 2 h at room temperature. Following PBS washes (3 × 10 min), slides were air-dried and mounted under a coverslip with a fluorescence mounting medium (Dako, # S3023).

Primary antibodies used were: goat anti-Phox2b (1:100; RD system, AF4940,), rabbit anti-peripherin (1:1000; Abcam,ab4666), guinea pig anti-Lmx1b (1:1000; (*63*)), goat anti-ChAT (1:100;Millipore, AB144p),), chicken anti-βGal (1:1000;Abcam, ab9361), chicken anti-GFP (1:1000;Aves Labs, GFP-1020), rabbit anti-Lmx1b (1:2000; (*63*)), guinea pig anti-Lbx1(1: 10000; (*63*), rabbit anti-DsRed (1:500;Takara bio, 632496), rabbit anti PKCdelta (1:1000;Abcam,ab182126,), rat anti-RFP(1:500; Rockland, 200-301-379).

All secondary antibodies were used at 1:500 dilution: donkey anti-chicken 488 (Jackson laboratories, 703-545-155), donkey anti-chicken Cy5 (Jackson laboratories, 703-176-155), donkey anti-goat Cy5 (Jackson laboratories, 705-606-147), donkey anti-rabbit 488 (Jackson laboratories, 711-545-152) donkey anti-rabbit Cy5 (Jackson laboratories, 712-165-153), donkey anti-rat Cy3 (Jackson laboratories, 711-495-152), and donkey anti-Guinea pig Cy3 (Jackson laboratories, 706-165-148). DAPI staining (Roth, #6335.1) was used at 1µg/mL to visualize cell nuclei in some experiments. Epifluorescence images were acquired with a NanoZoomer S210 digital slide scanner (Hamamatsu Photonics) and visualized with the NDPview2+ software; and confocal images with a Leica SP8 confocal microscope (Leica) with Leica Application suite X. Image formatting, including adjustments of brightness and contrast, and pseudo-coloring, was carried out in Adobe Photoshop (v.25.5.1) and FIJI.

## Behavioral experiments

### Timing and training

All behavioral experiments were conducted 3-4 weeks after viral injections and fiber optic implantation. One week after surgery, the mice were accustomed to handling, head fixation, and tethering to the patch cable during daily 10-minute sessions. This consisted in placing them into a plastic cylinder (4cm diameter) mounted onto a small aluminum breadboard (450mm X 450mm X 12.7mm, ThorLabs; MB4545/M), and head-fixing them using a custom-made fast fixation system (Ymetry, Paris, France) via a skull-attached head fixation implant. Their forepaws rested on the cylinder edge and the head protruded out. Mice were given either a 15% sucrose solution or a piece of almond or pasta in the home cage at the end of each session to reduce neophobia to these substances. During the lapping sessions, mice were given a 15% sucrose solution via a lick port. Chewing epochs were initiated by bringing a piece of almond or a piece of raw pasta within reach of the mouse’s jaw until the mouse grasped it between the jaws and held it in its forepaws. To highlight the jaw silhouette during video acquisitions, mice were illuminated from below and from the sides with white LED lights.

### Optogenetics and lick-induced photostimulation

To activate Sup5*^Phox2b^* neurons that express StCochR, a surgically implanted fiber-optic cannula was connected to a 473-nm DPSS laser (CNI, Changchun, China) via a patch cord (200 μm, 0.39 NA) using a zirconia mating sleeve Thorlabs). Simultaneous bilateral silencing of Sup5*^Phox2b^*expressing WichR1 was achieved using bifurcated fiber bundles (THORLABS BFYL2LS01) connected to the same laser source via an SMA905 connector. Spike 2 software and a 1401 data acquisition unit (Cambridge Electronic Design) were used to control laser output. Single continuous light pulses of 50–1000 ms or trains of 100 ms pulses at 5 Hz were used. In all behavioral experiments, the minimal laser output required to elicit a response was used, which was measured to be approximately 5 mW at the fiber tip using a digital power meter (PM100USB, Thorlabs). The laser output was digitized at 1 kHz by a NI USB-6008 card (National Instruments) and acquired using Spike 2 (CED Spike 2 Data Acquisition & Analysis Software).

To modulate optical stimulation or inhibition of Sup5*^Phox2b^*in a controlled and systematic manner, the timing of laser output was programmed (in Spike 2 software) such that the laser would be triggered mid-bout with a fixed delay after lapping onset. This delay was set to 480 ms, just under the average duration of lick bouts that we empirically defined as 4 successive licks (570ms) or more.

### Photometry

Single site fiber photometry recordings were made using a Doric fiber photometry system (Doric Lenses Inc, Canada) with 405 nm(isosbestic) and 465 nm excitation using Doric Neuroscience Studio software. Data were acquired at 12kHz.

### Automated markerless pose estimation

Spontaneous and light-evoked oromotor movements were filmed at portrait and profile angles with a CMOS camera (Jai GO-2400-C-USB) synchronized by a 5 V TTL pulse. The acquired frames had a resolution of 800 × 800 pixels and were streamed to a hard disk using 2ndlook software (IO Industries). The frames were then compressed using an MPEG-4 codec at a rate of 120 fps.

A ResNet-50 deep learning model was trained using DeepLabCut (version 2.3.8) on profile frames of mouse faces, to identify the genu of the jaw. This model was used to generate pose estimation of jaw in the experimental videos.

## Data analysis

### Fiber photometry

Custom-written Python scripts (Python version 3.7, Python Software Foundation) or Matlab code (MATLAB R2024a) were used to analyze behavioral and fiber photometry data. To match the acquisition rate of video recordings, fiber photometry and photostimulation data were resampled to 120 Hz. Photometry data was processed by first applying a low-pass filter (Butterworth) to the calcium-dependent 465 nm and isosbestic 405 nm signals with a 20 Hz cut-off. The 465 nm signal was then normalized using the function ΔF/F = (F-F0)/F0, where F is the 465 nm signal, and F0 is the least-squared mean fit of the 405 nm signal.

### Normalization of jaw and tongue pose estimation

Adjustments to the estimated positions of the jaw in Cartesian pixels were done by calibration against a 5mm scale bar within the video frame and the data was smoothed with a Savitzky-Golay filter. During optogenetic experiments, the position of the jaw was standardized to its average location 50-100ms before stimulation. For fiber photometry experiments, the jaw’s location was normalized to its highest vertical position (set as the resting position) during a recording session.

### Annotation of photometry data during ingestion

To detect biting versus chewing events during almond or pasta consumption, behavioral videos were annotated by eye using QuickTime Player version 7.7.9 to extract the start and end frames for each of these two behaviors within an 18-second interval. Biting events were characterized as instances when the mouse bit or gnawed the food material to break it into smaller pieces. Chewing events, on the other hand, were defined as the interval between two consecutive biting events, during which the mouse grinded the food material between its teeth before swallowing. The resampled and annotated video data was then aligned to the pre-processed photometry data in MATLAB to examine the calcium dynamics during biting and chewing of almond and pasta.

## Acknowledgments

We thank A. Delecourt and G. Firmin and the animal facility of IBENS for managing the mouse colonies, Astou Tangara and the imaging facility of IBENS, Carmen Birchmeier for the generous gift of the *Olig3:Cre* mouse and the anti-Lmx1b and anti-Lbx1 antibodies, Ofer Yizhar for the AAV1-EF1a-DIOstCoChR-P2A-mScarlet vector and Rémi Fournel for advice on data analysis and curation.

## Funding

INSERM: JFB, SS, ZC, CM

CNRS: JFB, SS, ZC, CM

ANR-19-CE16-0029-177 01, JFB

ANR-17-CE16-0006-01, JFB

FRM EQU202003010297, JFB

APP2001128 from the National Health & Medical Research Council, BD, AE.

## Author contributions

Conceptualization: JFB, BD, SS

Management of mice: ZC

Virus injections: SS, CMH

Immunohistology and behavioral tests: SS, JFB

Tracing experiments: SS

Data analysis SS, BD, AE, PB

Funding acquisition: JFB

Supervision: JFB, BD

Writing—original draft: JFB

Writing—review & editing: SS, BD

## Data and materials availability

“All data are available in the main text or the supplementary materials.”

**This PDF file includes**:

Figs. S1 to S#4

**Other Supplementary Materials for this manuscript include the following:**

Movies S#1 to S#8

**Sup. Fig.1:**
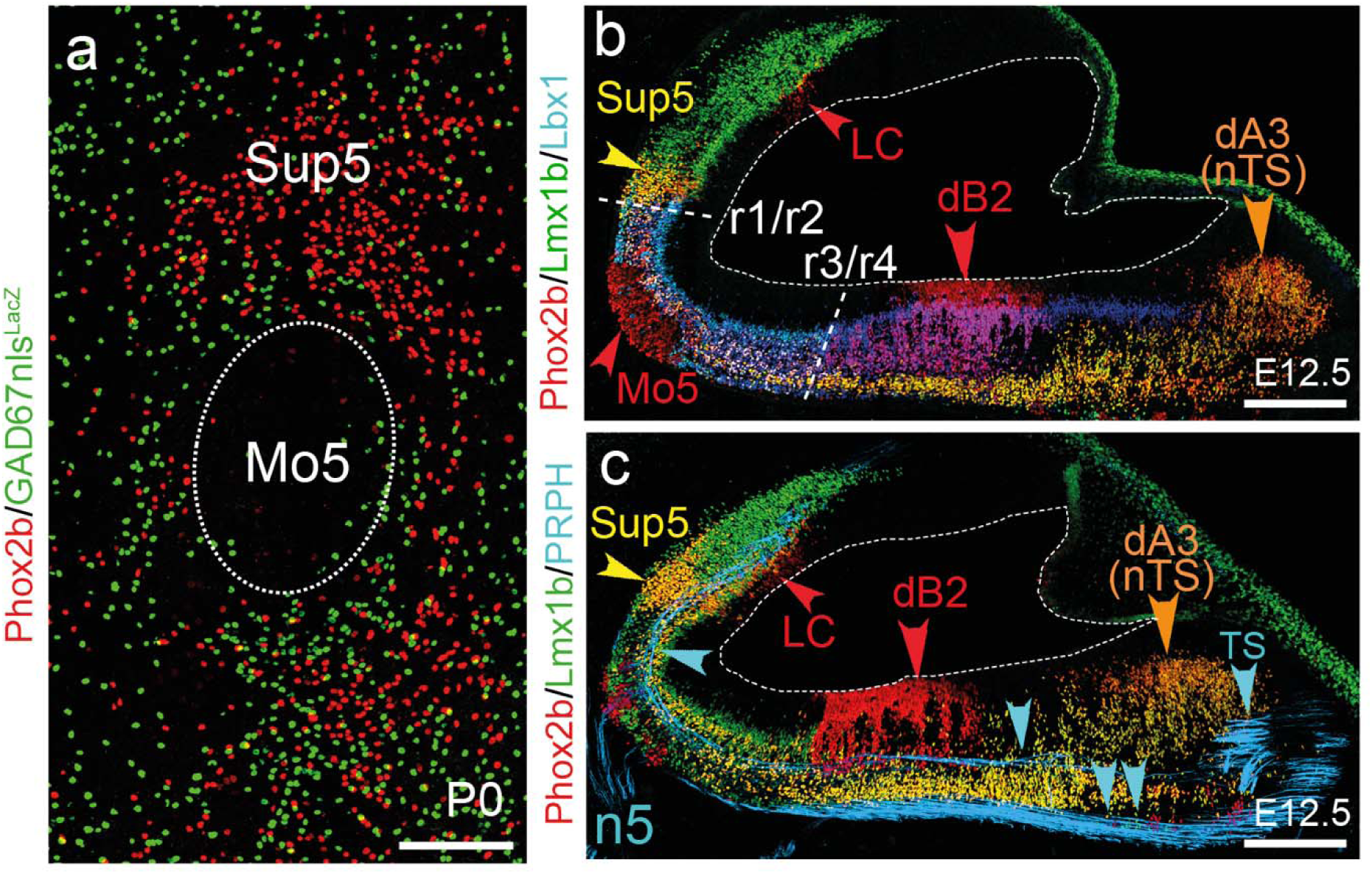
Genetic markers and ontology of Sup5*^Phox2b^*. (**a**) Section through Mo5 and Sup5*^Phox2b^* at P0 stained for the indicated markers. (**b**,**c**) Parasagittal sections of the brainstem at E12.5, stained with the indicated markers. The rhomboidal shape of the hindbrain entails that sagittal sections cut through different dorsoventral progenitor domains or their progeny (such as dB2 or dA3) at different rostro-caudal levels. (**b**) Sup5*^Phox2b^* develops just rostral to the r1/r2 boundary (stippled line), marked by the rostral limit of Lbx1. The r3/r4 boundary (stippled line) is the caudal limit of Lmx1b expression in Phox2b^+^ /Lbx1^+^ dB2 progeny (triple-labeled cells, white), caudal to which they are Phox2b^+^ / Lbx1^+^ / Lmx1b ^—^ (purple). (**c**) The emerging Sup5*^Phox2b^* lies close to the Me5 tract, which runs all the way to the caudal hindbrain (blue arrow). LC, locus coeruleus; n5, trigeminal nerve; nTS, nucleus of the solitary tract; PRPH, peripherin; TS solitary tract. Double blue arrow: spinal trigeminal tract; Sup5: Sup5*^Phox2b^*. The fourth ventricle is outlined with a stippled line.

**Supplementary Fig. 2.**
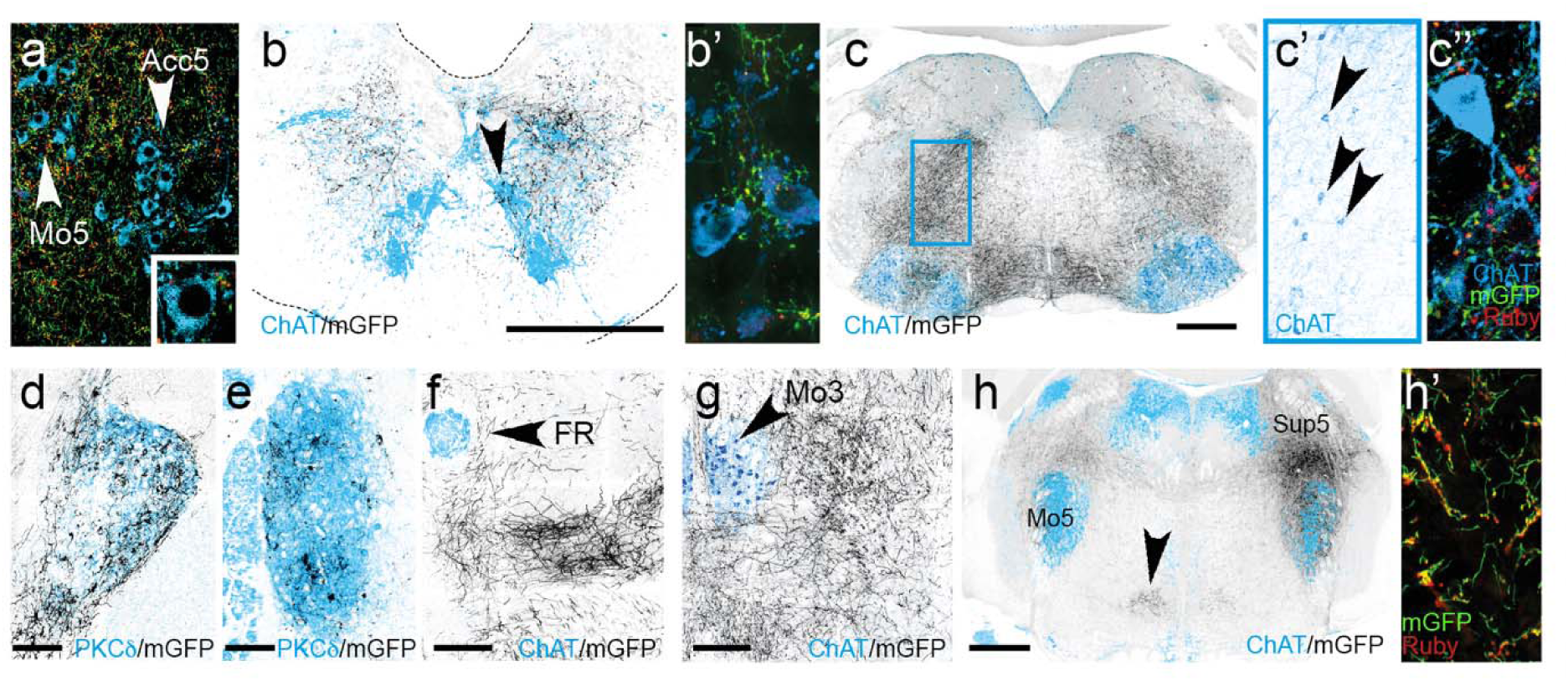
Additional sites of projections of Sup5*^Phox2b^*. (**a**) Section through Mo5 and Acc5 showing mGFP+ fibers and syp-Ruby+ boutons from Sup5*^Phox2b^* on both nuclei. Inset: close up of a neuron in Acc5. (**b**,**b’**) Projections of Sup5*^Phox2b^* on MoC (arrowhead in (**b**), high magnification in (**b’**). Blue: chAT; green: mGFP; red:syp-Ruby. (**c**,**c’**,**c’’**) The inferior salivatory nucleus located in the ventrolateral reticular formation receives projections (grey) on both sides (**c**), with ipsilateral predominance. (**c’**) Magnification of the blue boxed region in (**c**), showing the characteristically dispersed ChAT+ cells of the inferior salivatory nucleus in the reticular formation above Mo7 (arrowheads). (**c’’**) Close-up showing a ChAT+ salivatory nucleus cell covered with syp-Ruby boutons. (**d**-**g**) More rostral regions targeted by the Sup5*^Phox2b^*include the extended amygdala (Central amygdala (**d**) and the bed nucleus of the stria terminalis (**e**)); the ventral posteromedial thalamus (**f**) and the deep mesencephalic nucleus surrounding Mo3 (**g**). (**h**,**h’**) The caudal pontine reticular nucleus (arrowhead) located at the rostrocaudal level of Mo5 and Sup5 (h) receives dense inputs from the Sup5*^Phox2b^*. (**h’**) Higher magnification showing Syp-Ruby puncta in the pontine reticular nucleus. BNST: bed nucleus of the stria terminalis; CeA: central amygdala; DPME: Deep mesencephalic nucleus; FR: Fasciculus Retroflexus; Mo3: oculomotor nucleus, PnC: Pontine reticular nucleus caudalis; VPM: ventral posteromedial thalamus. Scale bars, **a, b, c** 500 µm; **d, e** 200 µm; **f, g** 100 µm.

**Supplementary Fig. 3:**
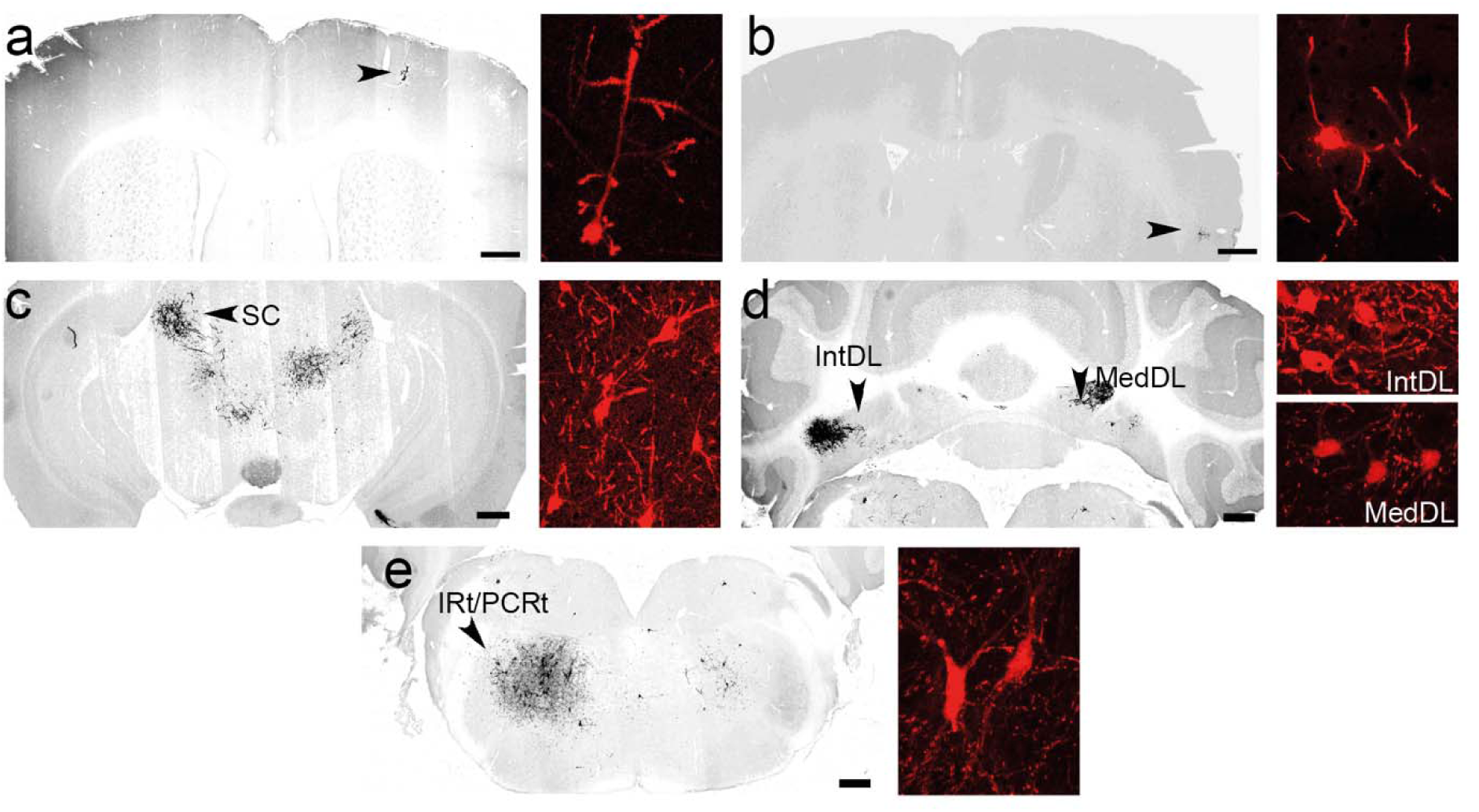
Additional sites of inputs to Sup5*^Phox2b^*. Inputs (mCherry+) are shown at low magnification (grey, left panels) and high magnification (red, right panels). (**a**,**b**), motor (**a**) and insular (**b**) cortices (Cx) provide contralateral input; (**c**) superior colliculus (SC) — more specifically its intermediate and deep lateral (motor-related) layers — projects ipsilaterally to Sup5*^Phox2b^*; (**d**) in addition to Lat (**Fig.3**) several deep cerebellar nuclei target Sup5*^Phox2b^*, including the intermediate dorsolateral DCN (IntDL) (ipsilaterally) and medial dorsolateral DCN (MedDL) (contralaterally); (**e**) Ipsilateral intermediate (IRt) and parvocellular (PCRt) reticular formations. Scale bars: **a**-**e**, 500 µm.

**Supplementary Figure 4:**
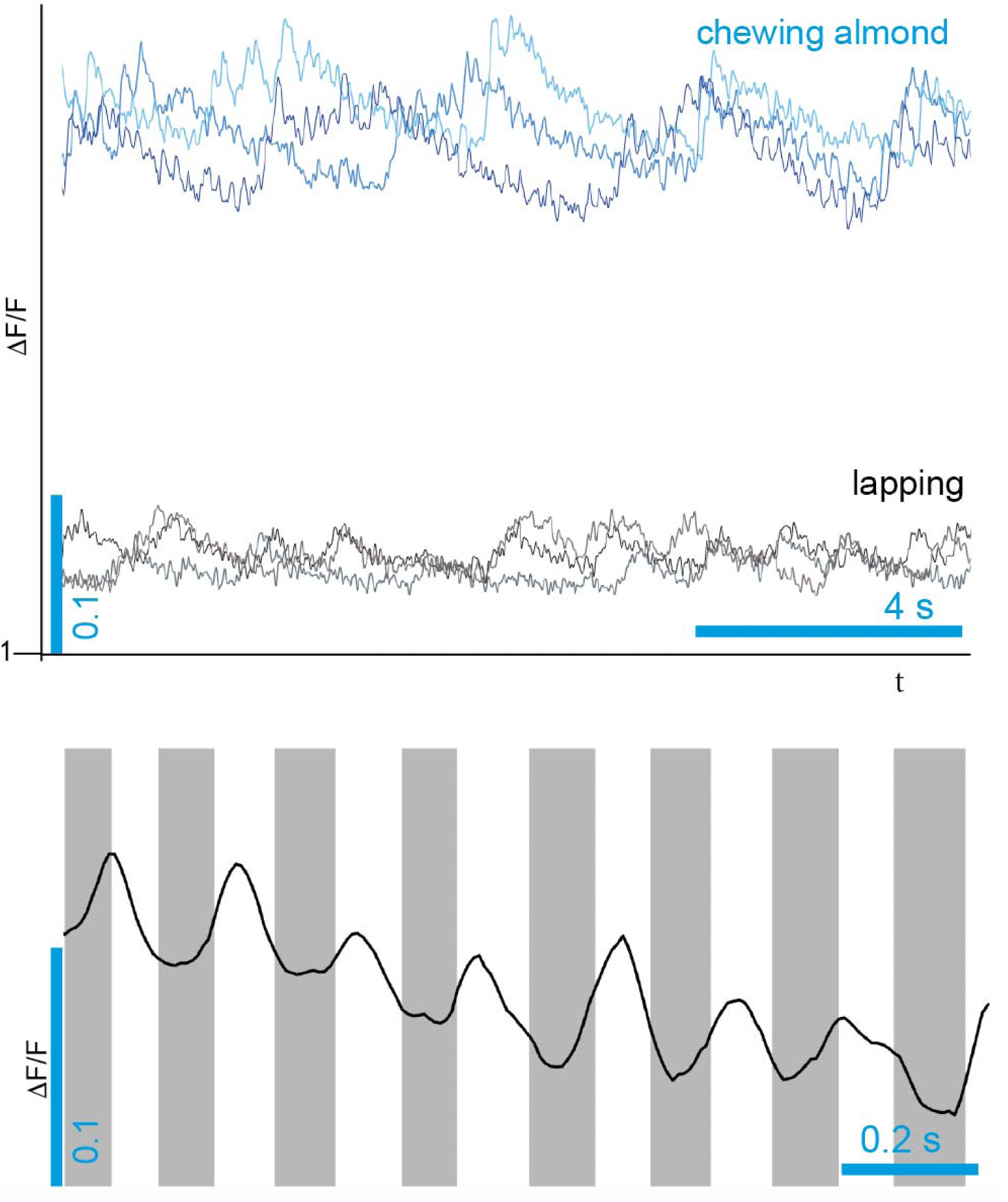
DF/F during lapping and chewing. **a**) DF/F comparison between lapping (3 lower traces) and chewing almond (3 upper traces) in 3 trials on one mouse. (**b**) DF/F while chewing an almond: increases in fluorescence precede each closure of the jaw (grey area).

**Movie S1**.

1000 ms optogenetic activation of Sup5^Phox2b^ at rest

**Movie S2**.

5 times 100 ms optogenetic activation of Sup5^Phox2b^ at rest

**Movie S3**.

1000 ms optogenetic activation of Sup5^Phox2b^ while licking

**Movie S4**.

1000 ms optogenetic activation of Sup5^Phox2b^ while chewing

**Movie S5**.

1000 ms optogenetic inhibition of Sup5^Phox2b^ at rest

**Movie S6**.

1000 ms optogenetic inhibition of Sup5^Phox2b^ while licking

**Movie S7**.

1000 ms optogenetic inhibition of Sup5Phox2b while chewing

**Movie S8**.

Synchronized DF/F trace and film of mouse biting and chewing an almond.

